# Chromatin profiling identifies putative dual roles for H3K27me3 in regulating transposons and cell type-specific genes in choanoflagellates

**DOI:** 10.1101/2024.05.28.596151

**Authors:** James M. Gahan, Lily W. Helfrich, Laura A. Wetzel, Natarajan V. Bhanu, Zuo-Fei Yuan, Benjamin A. Garcia, Rob Klose, David S. Booth

## Abstract

Gene expression is tightly controlled during animal development to allow the formation of specialized cell types. Our understanding of how animals evolved this exquisite regulatory control remains elusive, but evidence suggests that changes in chromatin-based mechanisms may have contributed. To investigate this possibility, here we examine chromatin-based gene regulatory features in the closest relatives of animals, choanoflagellates. Using *Salpingoeca rosetta* as a model system, we examined chromatin accessibility and histone modifications at the genome scale and compared these features to gene expression. We first observed that accessible regions of chromatin are primarily associated with gene promoters and found no evidence of distal gene regulatory elements resembling the enhancers that animals deploy to regulate developmental gene expression. Remarkably, a histone modification deposited by polycomb repressive complex 2, histone H3 lysine 27 trimethylation (H3K27me3), appeared to function similarly in *S. rosetta* to its role in animals, because this modification decorated genes with cell type-specific expression. Additionally, H3K27me3 marked transposons, retaining what appears to be an ancestral role in regulating these elements. We further uncovered a putative new bivalent chromatin state at cell type-specific genes that consists of H3K27me3 and histone H3 lysine 4 mono-methylation (H3K4me1). Together, our discoveries support the scenario that gene-associated histone modification states that underpin development emerged before the evolution of animal multicellularity.

## Introduction

Animal development depends on the ability to precisely regulate gene expression in time and space to specify cell types with different functions as well as the ability to maintain those cell identities after differentiation. Although the mechanisms underlying these processes have been extensively studied within animals^1,2^, their evolution is less clear due to the lack of information in the closest relatives of animals, the unicellular holozoans. A mechanistic account of gene regulation from diverse representatives within these groups will help clarify how animal developmental gene regulation evolved, which aspects of this process predate animals, and which emerged concomitant with animal evolution.

The regulation of genes during animal development occurs at all stages of gene expression with transcriptional regulation primarily driving differences between cell types. At the level of DNA sequence, animals not only rely on promoter proximal cis-regulatory sequences to regulate transcription but also utilize distal elements called enhancers^3,4^. Enhancers are cis-acting regulatory elements that can act over long distances (at least in bilaterians) to regulate target gene expression^4^. Enhancers have been identified in most major animal groups and are likely an ancestral component of the animal gene regulation toolkit^5-8^. Studies on the filasterean *Capsaspora owczarzaki*, however, failed to identify any major contribution of distal enhancer-like gene regulation in this group, suggesting they emerged only in the animal lineage ^9^. Whether enhancers are truly animal-specific, however, is still not clear as more sampling within unicellular holozoans is required. In addition, a clear definition of enhancers and what would constitute long-range activity in basal-branching metazoans and unicellular holozoans is lacking as is clear functional evidence for enhancer activity (reviewed in ^10^).

The regulation of chromatin and in particular histone post-translation modifications (hPTMs) is also key to animal development^11-15^. hPTMs are used in a variety of ways to regulate the expression of genes^15,16^. PTMs have generally conserved genomic distributions across eukaryotes, e.g., Histone H3 lysine 4 tri-methylation and lysine 27 acetylation (H3K4me3 and H3K27ac) are found at active genomic regions like promoters or active enhancers in diverse organisms^17-22^. Other modifications like H3K4me1 have varying distributions. In animals H3K4me1 is associated with enhancer activity and the relative levels of H3K4me3 and H3K4me1 have been used to distinguish promoters versus enhancers^23,24^. In *Arabidopsis thaliana*, H3K4me1 is found on the bodies of active genes^17^ while in the green algae, *Chlamydomonas reinhardtii*, H3K4me1 is broadly distributed throughout the genome but excluded from active regions^25^ and may play a role in silencing^26^.

While the capacity to turn genes on when they are required is central to cell-type specification, it is equally important to ensure that genes which should be inactive are maintained in this state and the ability to stably repress genes is essential for cell differentiation. Two prominent hPTMs associated with repression are methylation of H3K9 and H3K27^27^. H3K27me3 is deposited by Polycomb Repressive Complex 2 (PRC2)^28,29^ and is a key regulator of developmental genes in animals^29-32^. Interestingly, in plants Polycomb plays analogous roles but this may have evolved independently as there are several mechanistic differences and many of the key players in animals are not found in plants^33-36^. Additionally, recent work found a role for PRC2/H3K27me3 in transposon repression in diverse eukaryotes^37-51^ leading to the hypothesis that this is an ancestral role for PRC2^39^. Despite their importance, repressive PTMs have thus far not been studied in unicellular holozoans. In the filasterean *C. owczarzaki*, where active PTMS are well characterized, both the H3K9 and H3K27 methylation machinery have been lost^9^.

The choanoflagellates are the sister group to animals and therefore a key group for understanding early animal evolution^52-54^ (Fig. 1A). Choanoflagellates are a genetically diverse clade of unicellular heterotrophs which are found in virtually all aquatic environments^55^. The species *Salpingoeca rosetta* has emerged in recent years as a leading research organism due to several factors^52,56^. Firstly, *S. rosetta* has a dynamic life cycle with numerous cell types^57-59^ (Fig. 1B). These include two swimming cell types: slow swimmers which are the default state in high nutrient conditions and fast swimmers which are a dispersal stage induced in low nutrients. Their life cycle also includes a facultative multicellular state known as a rosette colony which is induced via specific environmental cues^60,61^. Recent work has shown that the formation of rosettes does not involve major transcriptional changes^62^. Instead, however, another cell type called thecate cells are very transcriptionally distinct from all swimming cell types providing a model for cell type-specific gene regulation. Thecate cells are physically attached to the substrate and are likely the diploid progeny of mating between haploid swimming cells^63,64^. The *S. rosetta* genome has been sequenced^58^ and the availability of functional tools including transfection^65^ and CRISPR-Cas9 genome editing^66^ make it a powerful system to dissect pre-metazoan gene-regulatory capabilities.

**Figure 1.**
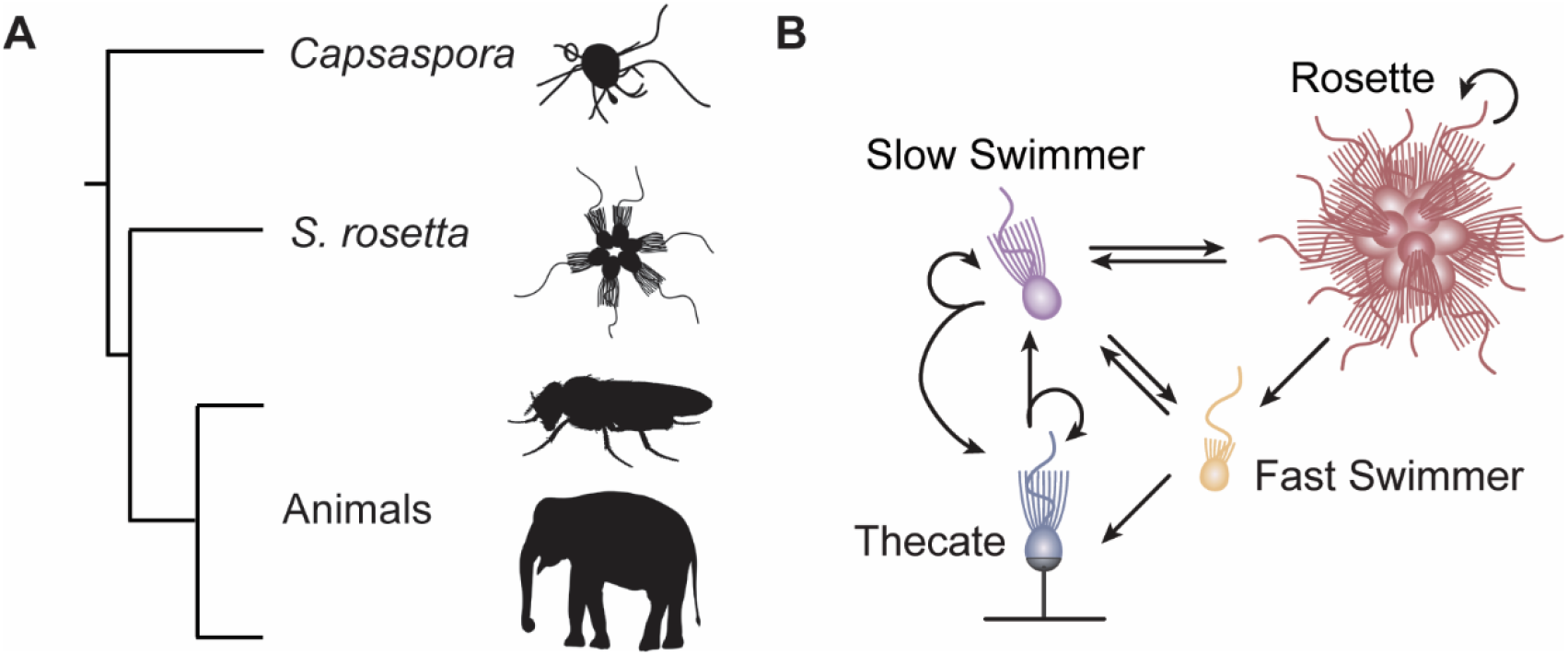
*Salpingoeca rosetta* as a model for pre-animal cell differentiation. (**A**) Phylogenetic tree highlighting the position of choanoflagellates (e.g., *S. rosetta*) as the sister group to animals. The filastereans (e.g., *Capsaspora owczarzaki*) are the sister group to animals and choanoflagellates. (**B**) The life cycle of *S. rosetta* consists of several distinct cell types representing different life history stages including single swimming cells (both slow swimmers and fast swimmers), a facultative multicellular stage called a rosette colony and a substrate-attached cell type called a thecate cell.

Despite their important phylogenetic position, there is currently little knowledge on gene regulation or chromatin in any choanoflagellate species. The role of one transcription factor, cRFX, in ciliogeneis has been reported^67^. To overcome this, we have mapped chromatin accessibility in *S. rosetta* using ATAC-seq and show that the majority of regulatory elements are in close proximity to transcriptional start sites (TSS). We have also catalogued the complement of hPTMs and described the genome-wide localization of several key hPTMs. This revealed a conserved pattern of hPTMs surrounding active genes while cell type-specific genes are occupied by H3K27me3-marked nucleosomes. H3K27me3 is also found on a subset of retrotransposons indicating a dual role for this modification in both transposon repression and cell type-specific gene regulation. Finally, we show a new putative bivalent state demarcating cell type specific genes when they are repressed.

## Results

### Gene regulation predominantly relies on promoter-proximal elements in *S. rosetta*

In all eukaryotes, the core promoter of genes drives gene expression and can mediate complex environmental sensing and differentiation^2,3^. Distal elements, like enhancers that impact transcriptional regulation of a gene add another layer of transcriptional regulation in animals. To understand what features of the cis-regulatory landscape in *S. rosetta* are shared with animals, their closest relatives, or other holozoans that lack distal enhancers, we employed ATAC-seq to map accessible chromatin regions genome-wide across cell types (Fig. S1A, B). Visualization of mapped reads showed clear peaks throughout the genome and high enrichment at TSSs (Fig. 2A, B and S1C). ATAC-seq signal showed a clear correlation between TSS enrichment and transcript levels (Fig. 2C). The majority of called peaks were in promoter regions or overlapped with predicted TSSs with a very small number of peaks distal to the TSS (Fig. S1D/E) which was also evident from manual inspection of the reads (Fig. 2A). A more detailed analysis revealed that approximately 75% of peaks directly overlapped a predicted TSS while more than 80% had their midpoint with a distance of -500 to +100 bps from a predicted TSS (Fig. 1D). Taken together, these data show that most cis-regulatory elements in *S. rosetta* are TSS-proximal with few distal regulatory elements.

**Figure 2.**
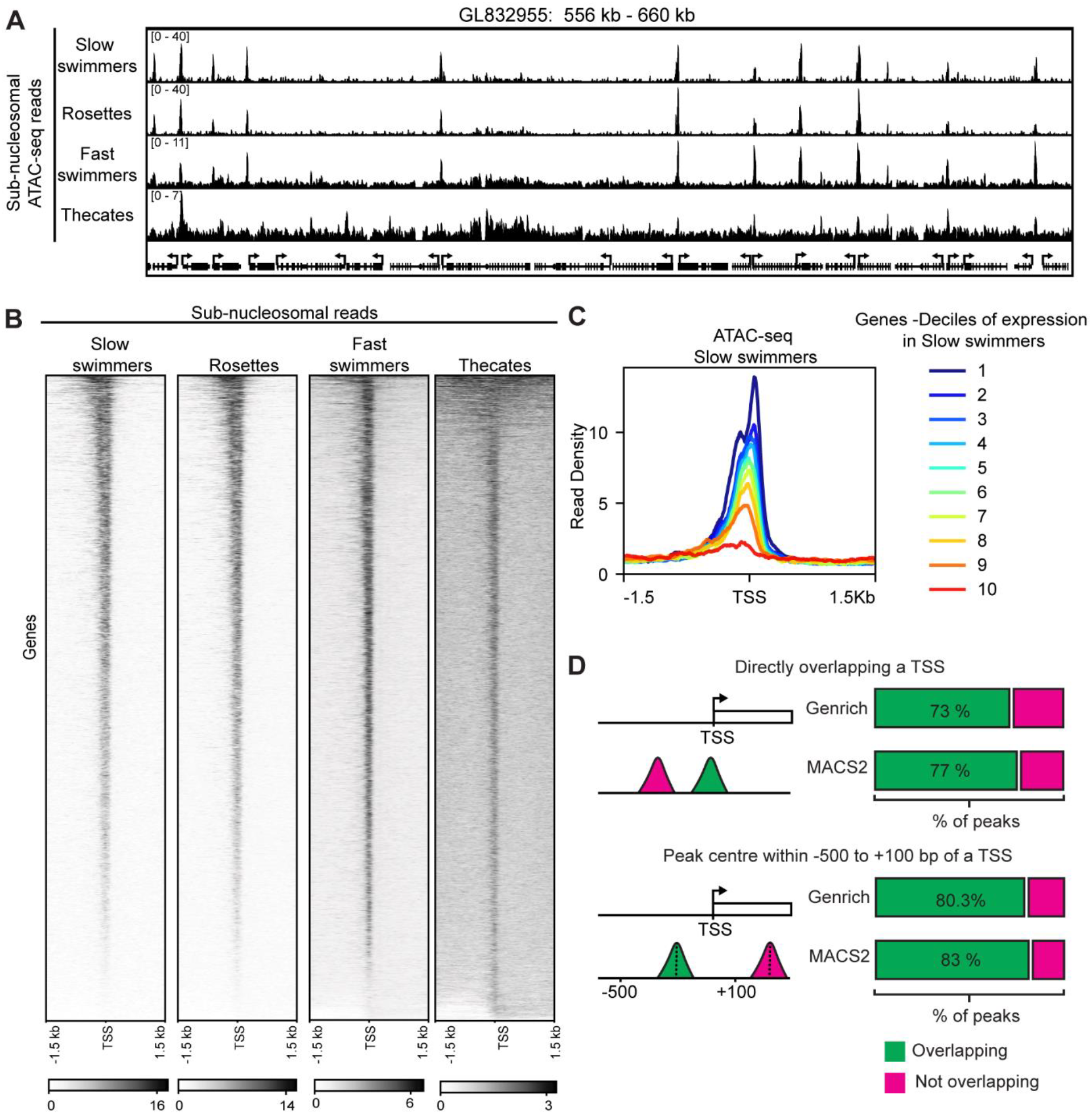
ATAC-seq on different cell types in *S. rosetta*. (**A**) Genome browser snapshot of ATAC-seq in the different *S. rosetta* cell types. Only sub-nucleosomal reads are shown. The cell type is shown on the left and the contig and region are shown on top. Annotated genes are shown along the bottom. (**B**) Heatmap of sub-nucleosomal ATAC-seq read density surrounding the TSS of all annotated genes (*n* = 11731). Cell type is shown on top. (**C**) Metaplot of ATAC-seq reads from slow swimmers surrounding the TSS for genes divided into ten deciles based on RNA expression (i.e. RPKM values) in swimming cells with 1 being the highest expressed and 10 the lowest. (**D**) Quantification of peaks which either overlap with predicted TSSs or with a midpoint located in a -500 to +100 bp window surrounding predicted TSSs. Peaks were called with either MACS 2 or Genrich.

### Genome-wide profiling of hPTMs reveals signatures for active and repressed genes

Given the prominent role of hPTMs in animal developmental gene regulation we next sought to understand the hPTM landscape in *S. rosetta*. Having defined the histones present (Fig S2A/B and Supplementary Table 1) we perform quantitative histone mass-spectrometry to identify hPTMs on samples extracted from both slow swimmers and thecate cells. We identified methylated and acetylated lysine residues in SrH3.1 and SrH4 (Fig. S2C). *S. rosetta* has all the common histone lysine methylation and acetylation sites seen in other eukaryotes, e.g., methylation of H3K4 and H3K36 and acetylation of H3K27 and H4K16. Importantly, we also identified both H3K9 and H3K27 methylation showing *S. rosetta* likely has both classical facultative and constitutive heterochromatin. We next mapped the genome-wide localization of 4 modifications: H3K4me3 and H3K27ac, two PTMs present at active regulatory elements genome-wide across eukaryotes, H3K4me1 which has variable distributions in different organisms but is generally present at both promoters and enhancers in animals and the PRC2-dependent modification H3K27me3. We optimized a native chromatin immunoprecipitation with sequencing (ChIP-seq) protocol using micrococcal nuclease (MNase) digested chromatin and performed ChIP-seq for the 4 hPTMs in duplicate in slow swimmers (Fig. S3A-D). K-means clustering of the distribution of PTMs around the TSS of all genes identified 5 broad groups of genes (Fig. 3A/B). Three clusters (Clusters 1,3 and 4) are associated with high levels of H3K4me1/3 and H3K27ac. A fourth cluster (Cluster 2) is depleted of these PTMs but was characterized by high levels of H3K27me3. The final cluster consists of genes with no obvious pattern (Cluster 5) which we excluded from further analysis. We then looked at the expression of genes from each cluster in both slow swimmers and thecate cells. RPKM values show that genes from clusters 1,3 and 4 are expressed in slow swimmers while cluster 2 genes show lower levels of expression (Fig. 3C). Interestingly, cluster 2 seems to contain genes that are upregulated in thecate cells while genes from the other clusters generally do not change in expression between cell types (Fig. 3C/D). Finally, analysis of ATAC-seq reads shows that the promoters of cluster 1,3 and 4 have an open, nucleosome-free region while cluster 2 genes do not (Fig. 3E and S3E). Inspection of genes from clusters 1, 3 and 4 revealed the major defining difference to be orientation within the genome (Fig. 3F). In each case, overlapping domains of H3K4me1/3 and H3K27ac are found flanking a nucleosome-free region at the TSS. In the case of cluster 4 genes these modifications are only seen downstream of the TSS but this may reflect the compact nature of the genome as an actively transcribed or H3K27me3-marked gene are always directly upstream (Fig 3A, B, F). Taken together this shows that hPTMs differentially demarcate active and inactive genes in *S. rosetta*.

**Figure 3.**
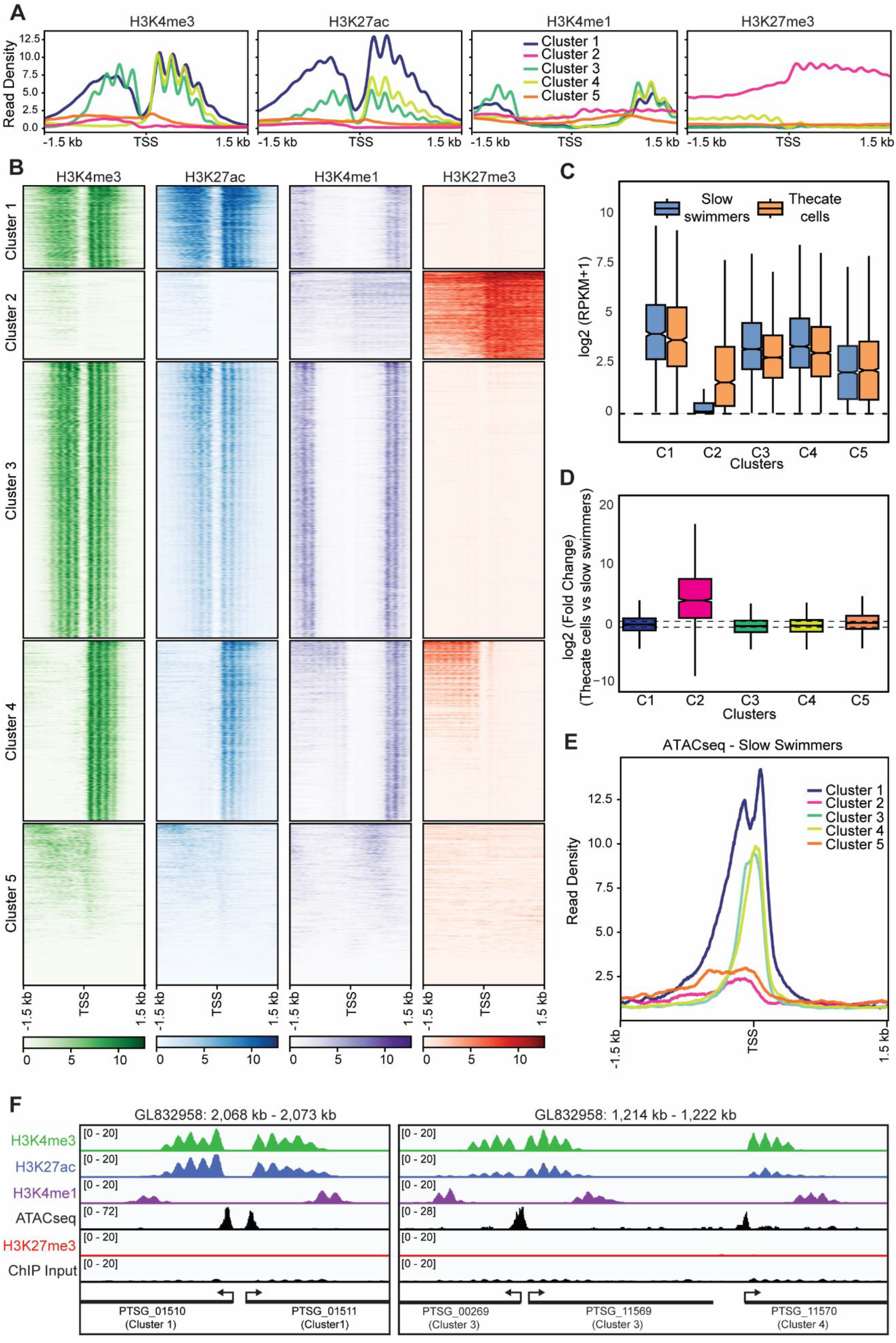
ChIP-seq of hPTMs reveals chromatin states of active and repressed genes. **(A)** Metaplot analysis of ChIP-seq with the indicated antibodies for each cluster of genes. Clustering was performed by k-means clustering. **(B)** Heatmap of data shown in A. (**C**) Box plot showing the log_2_ transformed RPKM values (+1) for genes in each cluster in slow swimmers and thecate cells. **(D**) Box plot showing the log_2_ fold change between Slow swimmers and Thecate cells for genes in each cluster. The dashed lines are at +1 and -1. The boxes show interquartile range, center line represents median, whiskers extend by 1.5× IQR or the most extreme point (whichever is closer to the median), while notches extend by 1.58× IQR/sqrt(n), giving a roughly 95% confidence interval for comparing medians. **(E)** Metaplot analysis showing ATAC-seq in swimming cells surrounding the TSS for genes from each cluster. **(F)** Genome browser snapshots of genes representative of clusters 1,3 and 4 showing ChIP-seq with the indicated antibodies along with input and ATAC-seq in slow swimmers. The genes are shown on the bottom as well as which cluster they belong to. The genomic scaffolds and positions are shown on top. Annotated genes are shown along the bottom. Cluster 1 (n= 1232), Cluster 2 (n= 1305), Cluster 3 (n= 4112), Cluster 4 (n= 2681), Cluster 5 (n= 2401).

### H3K27me3 and H3K4me1 co-occur on cell type-specific genes when they are repressed

Given the lack of knowledge on Polycomb-mediated repression in unicellular holozoans and its prominent role in animal developmental gene regulation, we decided to focus more closely on the cluster 2 genes which are strongly associated with the PRC2-mark H3K27me3. Looking at the expression of Cluster 2 genes we see that ∼75% are upregulated in thecate cells (Fig. 4A) but they only represent a small proportion of all thecate-upregulated genes (Fig. S4A). To understand what distinguished the Cluster2/upregulated genes from the other upregulated genes we first looked at the level of upregulation and see that the Cluster2 genes are generally very highly upregulated in thecate cells compared to others (Fig. S4B) and represent 86% of the top 500 upregulated genes (Fig. 4B). Further, in swimming cells, when these genes are H3K27me3 marked, we see that they are very lowly expressed or not expressed at all (Fig. 4C). Taken together, this shows that H3K27me3 marks a population of genes that are regulated in a cell-type-specific manner. We then looked closer at the pattern of hPTMs on these genes in slow swimmers (Fig. 4D/E). Unsurprisingly, they were marked by H3K27me3 which was the defining feature of cluster 2 from which they are derived. There were, however, two unexpected aspects of their hPTM signature. Firstly, H3K27me3 is very specific to the gene body of these genes rather than being localized on their promoters. Secondly, we see a surprising co-occurrence of H3K4me1 on these genes in a pattern which overlaps that of H3K27me3. Together, this shows that cell type-specific genes in *S. rosetta* are marked both by H3K27me3 and by H3K4me1 when repressed, a situation potentially analogous to bivalent chromatin in animals.

**Figure 4.**
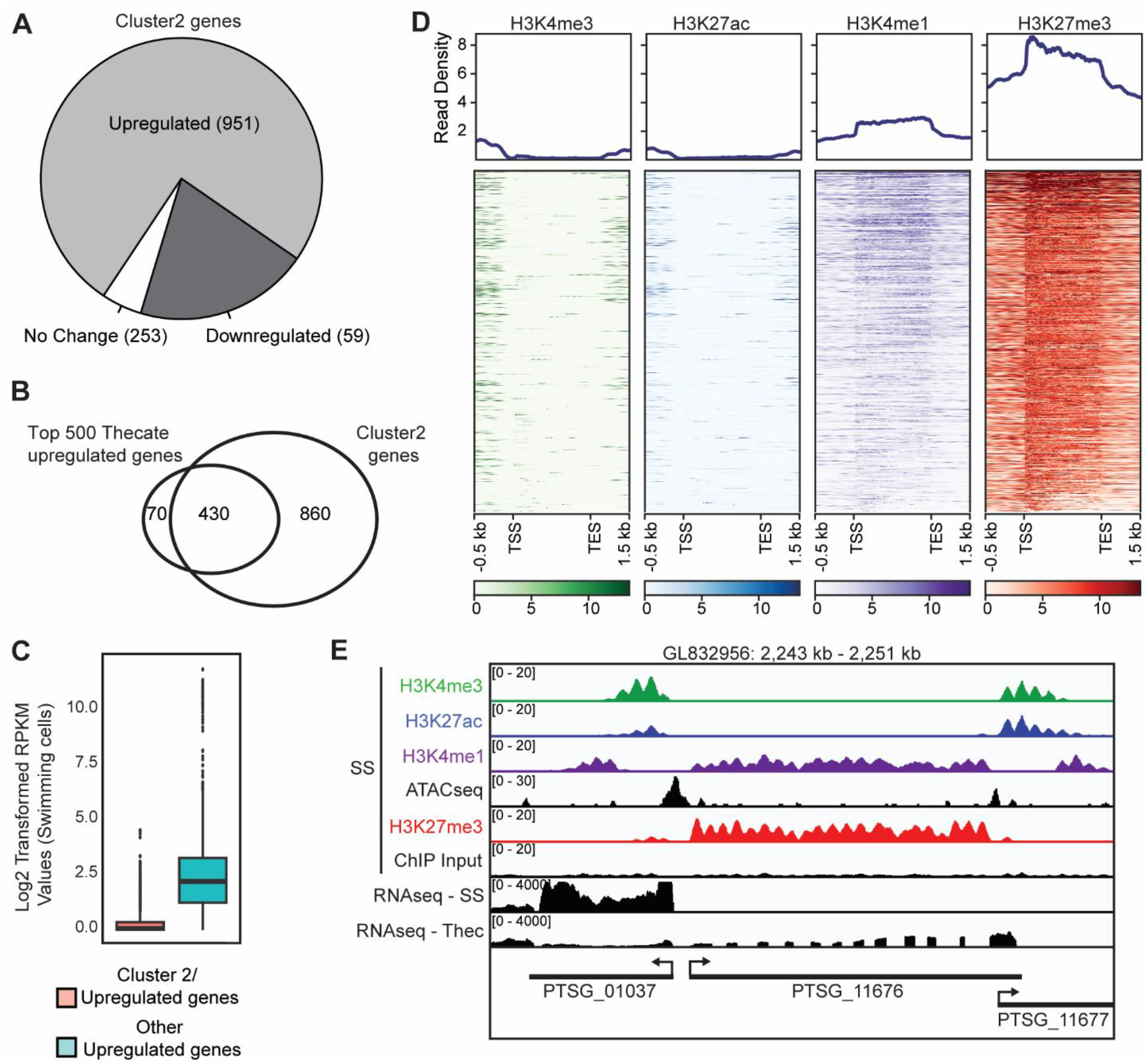
Thecate-specific genes are marked by H3K27me3 and H3K4me1 when silent. (**A**) Pie chart showing cluster 2 genes and whether they are upregulated, downregulated on unchanged between thecate cells and slow swimmers. Log_2_ fold change of +/- 1, as determined using DESeq2, was considered as a significant change. **(B)** Venn diagram showing the overlap between genes from cluster 2 and the top 500 genes upregulated in thecate cells vs slow swimmers. **(C)** Box plot showing log_2_ transformed RPKM values (+1) for genes in slow swimmers. Gene upregulated in Thecate cells are shown and separated based on whether they overlap with cluster 2 genes or not. The boxes show interquartile range, center line represents median, whiskers extend by 1.5× IQR or the most extreme point (whichever is closer to the median) and dots show outliers. **(D)** Heatmap and metaplot showing ChIP-seq with the indicated antibodies over cluster 2/thecate upregulated genes (n = 951). **(E)** Genome browser snapshot of a gene representative of Cluster 2 showing ChIP-seq with the indicated antibodies along with input, ATAC-seq in slow swimmers and RNAseq in both slow swimmers and thecate cells. The genomic scaffolds and positions are shown on top. Annotated genes are shown along the bottom. SS, Slow swimmers; Thec, Thecate cells.

### H3K27me3 labels LTR/Gypsy-like retrotransposons

Given the emerging roles of H3K27me3 in transposon repression in diverse eukaryotes, we wondered whether such a role may also be present in *S. rosetta*. A visual inspection revealed high levels of H3K27me3 at the end of many of the super-contigs in the genome. Given that many of the super-contigs are in fact full chromosomes^68^ we wondered if the H3K27me3 regions could be sub-telomeric regions. To address this, we looked at those super-contigs in the genome assembly where telomeric repeats are present within the sequence^58^. Of these, five out of six contained high levels of H3K27me3 enrichment adjacent to the telomeric repeats but no enrichment of the other modifications (Fig. 5A), while the remaining contig had a large gap in this region. In addition, we noticed that some H3K27me3-marked regions overlapped areas resembling transposons. To look further into this, we utilized RepeatMasker^69^ to annotate the genome using a previously published transposon annotation^70^. We then looked at hPTMs sitting on top of two prominent groups of transposons in the genome: LTR/gypsy-like retrotransposons and mutator-like element (MULE) DNA transposons. We observed enrichment of H3K27me3 but not the other modifications over LTR/gypsy-like retrotransposons but not MULE

**Figure 5.**
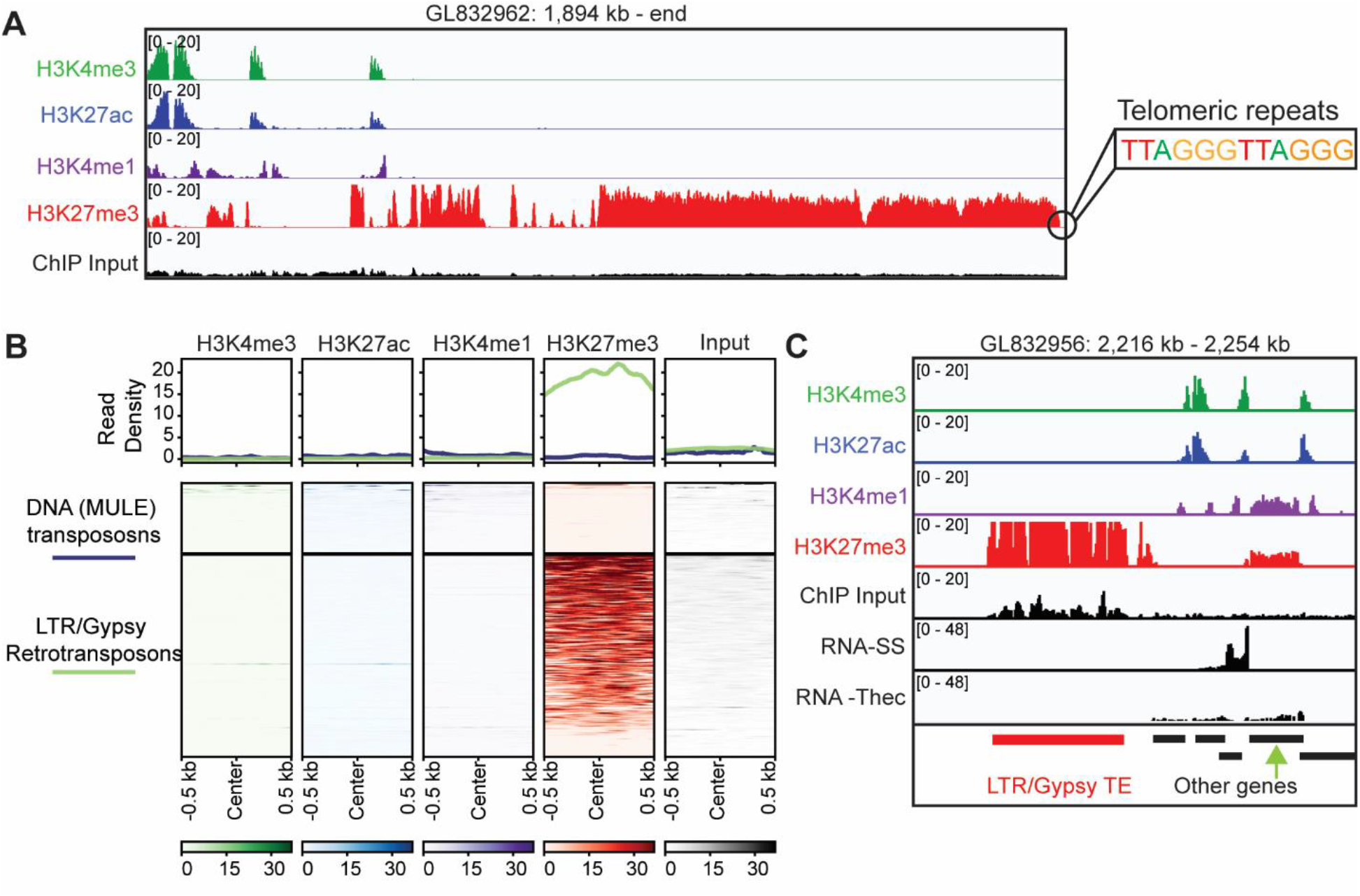
H3K27me3 labels sub-telomeric regions and a subset of transposable elements. **(A)** Genome browser snapshot of the sub-telomeric region of a chromosome showing ChIP-seq for the indicated antibodies and input. The genomic scaffold and position is shown on top. **(B)** Metaplot and heatmap showing ChIP-seq for the indicated antibodies and input over DNA(MULE) transposons (n=94) and LTR/Gypsy transposons (n=269). Reads here were not filtered to remove multi-mapping reads. **(C)** Genome browser snapshot of an example LTR/Gyspy retrotransposon showing ChIP-seq for the indicated antibodies and input along with RNAseq in slow simmers and thecate cells. The same region is shown in Fig. 5E, and the cluster 2 gene shown there is indicated here with a green arrow. The genomic scaffold and position is shown on top. Annotated transposons and genes are shown along the bottom. SS, Slow swimmers; Thec, Thecate cells.

DNA transposons (Fig. S4C) which is even more evident if we include all reads (i.e. not excluding muti-mapping reads) (Fig. 5B/C). The example in Fig. 5C is the same genomic region as in Fig. 5E but zoomed out. The cluster 2 gene shown in Fig. 4E is annotated with a green arrow. Comparing this with the annotated LTR/gypsy-like element highlights the co-occurrence of H3K4me1 on repressed cell type-specific genes but not the transposable elements.

## Discussion

Here we present the first analysis of chromatin accessibility and hPTMs in choanoflagellates and present findings that have broad implications for our understanding of gene regulation at the origin of animals. We observe no clear evidence for distal enhancers like those found in animals. Together with work in *C. owczarzaki*^9^, these data suggest that animals elaborated on core chromatin and transcription machinery from their holozoan ancestor to regulate gene expression with distal cis-regulatory elements. More mechanistic work is necessary to uncover the evolutionary changes that enabled distal enhancers to become a widespread feature of developmental regulation in animals. We cannot exclude the possibility that *S. rosetta* may have some features of enhancers that are difficult to disentangle in a condensed genome with a mean intergenic distance of 885 bp^71^. For example, the core promoter from one gene may influence the expression of neighboring genes, similar to the original discovery of enhancers whereby an SV40 promoter enhanced the transcription of a distant beta-globin gene^72^. One alternative cause may be the small size of the genome (∼55Mb). Animals that have undergone massive genome size reductions sometimes have limited intergenic enhancers but often retain distal enhancer-type elements within introns^73-75^. We do not see any evidence for such intronic regulatory sequences, arguing against a loss of enhancers during a reduction of the genome size. Notably, transcriptomes from diverse choanoflagellate species suggest that some have a larger gene repertoire than the two model species, *S. rosetta* and *Monosiga brevicollis*, that have sequenced genomes. Additional genomes may therefore reveal other features of choanoflagellate chromatin biology and help reveal the origin and evolution of animal chromatin-based gene regulation.

The ability to stably repress genes in a cell type-specific manner is a crucial component of animal developmental gene regulation. This is achieved partly by the Polycomb-repressive system. Here, we present evidence for a dual role for PRC2 and its modification H3K27me3 in the regulation of cell type-specific genes and transposons in *S. rosetta*. This raises several important evolutionary considerations. Recent work has highlighted the previously underappreciated role of PRC2 in regulating transposons across diverse eukaryotes leading to a hypothesis that this may represent an ancestral role^39,45^. Our data are consistent with this, suggesting that this function was at least present in the last common ancestor of animals and choanoflagellates. The role of H3K27me3 in regulating cell type-specific genes is more complex. We show that H3K27me3 marks genes in a pattern that is consistent with a similar role as in animals. However, it may also represent an independent co-option in the *S. rosetta* lineage. There are therefore two possibilities: (1) H3K27me3 was already used in the last common ancestor of animals and choanoflagellates to regulate cell type-specific genes and this role was then inherited and retained in both lineages, or (2) H3K27me3 has been independently co-opted for such a function in both lineages. The mechanisms of PRC2 recruitment to chromatin are currently unknown in *S. rosetta*; therefore, further studies of those molecular mechanisms will likely better facilitate comparisons with animals to potentially resolve these evolutionary scenarios. In addition, the role of the other major Polycomb-repressive complex, PRC1, is not known in *S. rosetta* and is, in fact, not understood in any system outside of plants and bilaterian animals. It was recently shown by us and others that choanoflagellates contain a variant PRC1 complex, which is the evolutionary precursor of animal PRC1 complexes^36,76^. Future work on the role of this complex will be necessary to determine whether it co-operates with PRC2 in the regulation of transposons and/or cell type-specific genes in *S. rosetta*.

We show that in addition to being marked by H3K27me3, cell-type-specific genes are also marked by H3K4me1 when repressed. H3K4me1 may therefore have some putative roles at repressed gene in addition to its canonical roles as it is also found at active genes in a manner that matches what is seen in other systems. This situation is potentially analogous to “bivalent” chromatin domains in animals which are regions marked by both H3K4me3 and H3K27me3 ^77,78^. There are, however, several key differences. The bivalent state was first identified in embryonic stem cells^78,79^ and later in other animal species^80,81^ and is mostly associated with the co-occurrence of H3K27me3 and H3K4me3 on promoters^78,82-84^ and not H3K4me1. Some regions are, however, marked by H3K27me3 and H3K4me1 such as poised enhancers^77,84-89^. In *S. rosetta*, H3K27me3 and H3K4me1 co-occur over gene bodies rather than at regulatory elements, as in the cases outlined above. Despite these differences, it may be that there is a similar role.

Bivalent chromatin was originally believed to prime genes for rapid activation upon differentiation. However, this has recently been called into question, and H3K4me3 at bivalent regions is now thought to prevent DNA methylation and therefore prevent irreversible silencing^90-93^. It is therefore possible that H3K4me1 in *S. rosetta* also prevents some higher level of repression of cell type-specific genes and therefore bookmarks them for later activation. It is highly unlikely that this occurs through DNA methylation as the machinery for methylation is absent in *S. rosetta*^*94*^. It may be that this is preventing accumulation of other hPTMs. Outside of animals, a similar type of bivalent chromatin was recently described in the plant *Brassica napus* where it is associated with tissue-specific genes^95^. Further studies will be needed to elucidate the functional relevance of this putative bivalent state and how it may relate to bivalent promoters and/or poised enhancers in animals.

## Materials and Methods

### Choanoflagellate cultures and media preparation

All experiments were performed using *Salpingoeca rosetta* in co-culture with a single bacterial food source, *Echinicola pacifica* (ATCC PRA-390, strain designation: SrEpac)^63,96^.

Artificial sea water – Keller formula (AKSW), high nutrient media (HN) and cereal grass media (CG) were prepared as previously described^63,65,96^.

ATSM D 1141 Artificial Seawater (hereafter ASW) was sourced commercially (ASTM D 1141, Ricca Cat No. 8363-5). RA media was prepared from *Porphyra umbilicalis* as previously described ^61^. Briefly 10 g of dried algae was added to 1 L of ASW and incubated mixing for 1 hour at room temperature. This was then sterile filtered and RA media was prepared by diluting this 1:4 in ASW along and 1:100 dilutions of 1000x (Potassium Iodide, Sodium Nitrate, Sodium Phosphate), 1,000x L1 vitamins^97^, and 1,000x L1 trace metals^97^.

### ATAC-seq

Nuclei were isolated from different cell types: (1) slow swimmers, (2) rosettes, (3) fast swimmers, and (4) thecate cells with two independent replicates. Cultures containing single cell types were produced as described previously^62,67^. Slow swimmers were generated by maintaining cells in HN media. Rosettes were induced with outer membrane vesicles from *Algoriphagus machipongonensis* as previously described^98^. Fast swimmers were grown 3 days in HN and then heat shocked at 30°C for 2.75 h. Slow swimmers, rosettes, and fast swimmers were harvested by pelleting at 2400 x g for 5 minutes, washed with 50 mL AKSW, re-pelleted at 2400 x g for 5 minutes, counted with the Luna cell counter (Logos Biosystems), diluted to 50 million cells/mL, and 10 million cells were pelleted at 2700 x g. Thecate cells were derived from an isolate of SrEpac, called HD1^63^, and maintained in 10% CG in AKSW (vol/vol) in petri dishes. To harvest thecate cells, plates were washed with 16.7 mL of AKSW, cells lifted from the plate with a cell scraper, and filtered onto a 3 μm polycarbonate membrane filter to concentrate. Filtered cells were pelleted at 2700 x g for 5 minutes, washed with 50 mL AKSW two times, re-pelleted at 2700 x g for 5 min, counted with Luna cell counter (manufacturer …it’s in my genome editing paper), diluted to 50 million cells/mL, and 10 million cells pelleted at 2700 x g. All cell types were resuspended in 200 μL freshly prepared pretreatment buffer (10 mM citric acid, 100 mM Lithium Acetate, 10% (w/v) PEG 8000 pH 8.5 with Tris, 100 nM papain, and 10 mM thioglycolic acid) and incubated at room temperature for 22 minutes. Nuclei were isolated in four steps: wash, strip, lyse, and purify. To wash, cells were pelleted at 1200 x g for 5 minutes, the supernatant discarded, and resuspended in 200 μL of 0.7 M sorbitol in 1x PBS and 1% (w/v) BSA, and pelleted at 1200 x g for 5 minutes. Pellets were resuspended in 250 μL cold buffer L (10 mM HEPES-KOH pH 7.9, 0.2 mM MgCl_2_, 10 mM KCl, 0.1 mM EDTA-KOH pH 8.0, 0.5 mM EGTA-KOH pH 8.0, 0.5 mM DTT, 0.5 mM Pefabloc-SC, and 1x Roche Complete EDTA-free protease inhibitor cocktail) and incubated for 10 minutes on ice. To lyse cells, 0.05% IGEPAL CA-630 was added, cells were incubated on ice for 10 minutes, and then samples were passed through a 30G needle ten times. Lysed cells were pelleted at 1000 x g for 5 minutes at 4°C and the supernatant was removed. Pellets were resuspended in 250 μL buffer L with sucrose (Buffer L, 250 mM sucrose, 0.5 mM DTT, 0.5 mM pefabloc, and 1x Roche protease inhibitor solution), spun at 1000 x g for 5 minutes at 4°C, and both steps repeated.

For transposition, nuclei were pelleted at 1000 x g for 5 minutes at 4°C, resuspended in 25 μL of 2x TD buffer and 2.5 μl of Tn5 transposase from the Nextera DNA Library Prep kit (Illumina, San Diego, CA), and incubated at 37°C for 30 minutes. DNA was purified using the MinElute kit (Qiagen, 28004) per PCR purification protocol provided by the manufacturer. Transposed DNA was originally amplified and barcoded in a PCR reaction using NEBnext PCR master mix (NEB, M0544) and 1.25 μM forward and reverse primers originally described in ^99^, using the following PCR conditions: 72°C for 5 minutes; 98°C for 30 seconds; and thermocycling at 98°C for 10 seconds, 63°C for 30 seconds and 72°C for 1 minute. To reduce GC and size bias in the PCR, we monitored the PCR reaction using qPCR in order to stop amplification before saturation. To do this, we amplified the full libraries for five cycles, after which we took an aliquot of the PCR reaction and added 10 μl of PCR cocktail with Sybr Green at a final concentration of 0.6x. We ran this reaction for 20 cycles to determine the additional number of cycles needed for the remaining 45 μL reaction. The libraries were purified using a Qiagen PCR cleanup kit. Libraries were amplified for a total of 10-12 cycles. An additional 0.9X SPRI bead cleanup was performed to eliminate a contaminating 50 bp peak. Samples were pooled, quantified using qPCR and sequenced on an Illumina HiSeq 2500 to generate 50bp PE reads.

### ChIP-seq

For ChIP-seq, approximately 500 million cells were used per experiment. Cells were seeded at

∼10,000 per ml in RA and grown for 24 hours at 27°C. Cells were centrifuged at 2400 x g for 5 minutes at 4°C (all further centrifugation steps and washed used these parameters unless otherwise specified). They were then washed once with ASW and resuspended in ASW, this time combining them into one 50 ml tube and cell number was quantified. Following centrifugation, the pellet was resuspended in 5 ml ice-cold lysis buffer (50 mM HEPES pH 7.6, 100 mM NaCl, 0.5 mM MgCl_2_, 10% (v/v) glycerol, 1% (v/v) Triton x-100, 10 mM sodium butyrate, 2 mM DTT, 2 mM Pefabloc-SC, 2x Complete EDTA-free Protease inhibitor cocktail) and incubated on ice for 30 seconds. Released nuclei were collected by centrifugation and washed once in nuclease digestion buffer (10 mM Tris-HCl pH 8.0, 10 mM NaCl, 3 mM MgCl2, 0.1% (v/v) IGEPAL CA-630, 0.25 M sucrose, 3 mM CaCl_2_, 2 x Complete EDTA-free Protease inhibitor cocktail). The nuclei were then resuspended in 1 ml of nuclease digestion buffer per 500 million cells and placed in a dry bath at 37°C for 5 minutes. Micrococcal nuclease (MNase) (ThermoFisher Scientific, EN0181) was added to the warmed nuclei at a final concentration of 450 U/ml and cells were incubated for exactly 5 minutes at 37°C with gentle mixing by inversion every minute. Digestion was halted by addition of 8 μL 0.5 M EDTA pH 8.0 per ml on ice. The samples were then centrifuged, and the supernatant was kept and placed on ice (S1). The remaining pellet was resuspended in nucleosome release buffer (10 mM Tris-HCl pH 7.5, 10 mM NaCl, 0.2 mM EDTA, 2x Complete EDTA-free Protease inhibitor cocktail), placed rotating end over end at 4°C for 1 hour and passed 5 times through a 27 G needle. Following centrifugation, the supernatant (S2) was retained and then added to the S1 (nucleosome solution). Digestion to majority mononucleosomes was confirmed by extracting DNA from 50 μL followed by agarose gel electrophoresis.

For each ChIP experiment 100 μL of mononucleosomes per antibody (+100 μL for input) was diluted 10-fold with ChIP incubation buffer (70 mM NaCl, 10 mM Tris-HCl pH 7.5, 2 mM MgCl_2_, 2 mM EDTA, 0.1% Triton, 1 x Roche Complete EDTA-free Protease inhibitor cocktail). 1 ml aliquots were then made and the antibodies (Listed in Supplementary Table 2) added and incubated rotating at 4°C overnight. The next day, CaptivA® Protein A Affinity Resin (Repligen, CA-PRI-0025) (40 µl slurry per ChIP, hereafter called “beads”) was pre-blocked in ChIP incubation buffer supplemented with 1 mg/ml BSA and 1 mg/ml yeast tRNA, for 1 hour at 4°C. The beads were collected by centrifugation at 1000 x g for 1 minute at 4°C (all centrifugations and washes from this point were performed using these parameters) and washed three times in ChIP incubation buffer. After being aliquoted into 1 ml tubes, the beads were collected by centrifugation and the antibody-nucleosome solutions added. This was incubated rotating at 4°C for 1 hour before the beads were collected by centrifugation and washed 5 times in ChIP wash buffer (20 mM Tris-HCl pH 7.5, 2 mM EDTA, 125 mM NaCl, 0.1% (v/v) Triton X-100) with 5-minute incubations rotating at 4 °C between each wash. Following a final wash with TE buffer, the beads were resuspended in 100 μL fresh elution buffer (1 % (w/v) SDS, 0.1 M NaHCO_3_) and incubated shaking at room temperature for 30 minutes. The beads were then collected by centrifugation and the supernatant retained. ChIP DNA was purified using the ChIP DNA Clean and Concentrator kit (Zymo, D5206) using the manufacturer’s instructions and eluted in 10 µl.

ChIP libraries were generated by the QB3-Berkeley Functional Genomics Laboratory (FGL) and sequenced at Vincent J. Coates Genomics Sequencing Laboratory (GSL) at UC Berkeley. Samples were checked for concentration and size using the Qubit dsDNA High Sensitivity assay (ThermoFisher Scientific, Q32851) and Agilent Fragment Analyzer with the DNA High Sensitivity NGS assay (Aligent, 5067-4626). Subsequently, the input material was size-selected using a double-sided bead cleanup at a 0.55x / 1.8x bead ratio using Kapa Pure Beads. Library preparation was carried out using the KAPA HyperPrep kit for DNA (KK8504). Truncated universal stub adapters were used for ligation, and indexed primers were used during PCR amplification to complete the adapters and to enrich the libraries for adapter-ligated fragments. After PCR, the libraries were cleaned at 0.9x bead ratio to remove smaller insert fragments and dimers. Samples were then checked for quality on an AATI (now Agilent) Fragment Analyzer. Illumina sequencing library molarity was measured with quantitative PCR with the Kapa Biosystems Illumina Quant qPCR Kits on a BioRad CFX Connect thermal cycler. Libraries were then pooled evenly by molarity and sequenced on a shared Illumina NovaSeq6000 150PE S4 flowcell. Raw sequencing data was converted into fastq format sample specific files using the Illumina BCL Convert V4 software on the sequencing center’s local Linux server system.

### Histone extraction and mass spectrometry

For histone extraction, thecate cells or swimming cells were seeded at approximately 10,000 cells per ml in RA and grown for 24 hours. Approximately 200 million cells were used per replicate. Cells were centrifuged at 2400 x g for 5 minutes at 4°C (all further centrifugation steps and washed used these parameters unless otherwise specified) in 50 ml tubes. They were then washed once with AKSW and following centrifugation each tube was resuspended in 1 ml ASW. This was then carefully layered on top of 2ml percoll solution (160 μl percoll brought to 2 ml with ASW) in a 15 ml tube and centrifuged at 1000 x g for 10 minutes at 4°C with the brakes off. The supernatant was carefully removed, the pellets resuspended in ASW and combined into one 50 ml tube and cells were counted. Following centrifugation, cells were resuspended in 1 ml lysis buffer (see above) per 50 million cells and incubated on ice for 30 seconds. The nuclei were collected by centrifugation and directly resuspended in 800 μL ice-cold 0.4 N H_2_SO_4_ and left at 4°C for 2 hours. They were then centrifuged at 16,000 x g for 10 minutes at 4°C and the pellet discarded. The histones were precipitated by drop-wise addition of 264 μL ice-cold 100% TCA followed by overnight incubation at 4°C. Histones were pelleted by centrifugation at 16000 x g for 10 mins at 4°C followed by three washes with ice-cold acetone. The histone pellet was allowed to air dry for approximately 15 minutes at room temperature and resuspended in 50-100 μL of water. Histone concentrations were determined using Bradford assay with BSA as standard.

Sample preparation and mass spectrometry were performed as previously described ^100^. Propionic acid derivatization was carried out in 20ug of purified histones. About 4-19 amino acid-long peptides were generated using propionic anhydride derivatization, followed by digestion with 1 μg trypsin and desalting for bottom-up mass spectrometry. The peptides were then analyzed using a Thermo Scientific Acclaim PepMap 100 C18 HPLC Column (250mm length, 0.075mm I.D., Reversed Phase, 3 um particle size) fitted on an Vanquish™ Neo UHPLC System (Thermo Scientific, San Jose, Ca, USA) using the HPLC gradient: 2% to 45% solvent B (A = 0.1% formic acid; B = 95% MeCN, 0.1% formic acid) over 50 minutes, to 95% solvent B in 10 minutes, all at a flow-rate of 300 nL/min. The sample (5 μl of 1 μg/μl) was analyzed in a QExactive-Orbitrap mass spectrometer (Thermo Scientific) using data-independent acquisition (DIA). It consisted of full scan MS (m/z 295−1100) acquired in Orbitrap with a resolution of 70,000 and an AGC target of 1x106. Tandem MS was acquired in centroid mode in the ion trap using sequential isolation windows of 24 m/z, AGC target of 2x105, CID collision energy of 30 and maximum injection time of 50 msec.

### Data analysis

All analysis software used are listed in Supplementary Table 3.

### ATAC-seq data analysis

ATAC reads were initially processed using Trimmomatic^101^ to remove adaptors. They were then mapped to the genome using Bowtie2^102^ (with the “-very-sensitive” option) and converted to BAM files and sorted using SAMtools^103^. PCR duplicates were removed using Picard and low-quality reads were removed using SAMtools view (“-q 30” option). SAMtools was used to extract sub-nucleosome sizes reads, i.e. reads with insert sizes less than 100 bps. DeepTools^104^ was used to generate PCA and Pearson correlation plots. Replicates were merged together for visualization and BigWig files were generated using bamCoverage (--binsize 10 –effectiveGenomesize 55000000 –normalizeUsing RPGC) and visualized using the Integrative Genomics Viewer (IGV) ^105^. Heatmaps and profile plots were generated using deepTools.

Peak calling was performed either with Genrich (-j -y -r -v options) which takes both replicates as input or with MACS 2^106^ for individual samples (-f -BAMPE option --keep-dup all). Sub-nucleosomal reads were used as input for Genrich while all reads were used for MACS2. In the case of MACS2 consensus peaks sets for each cell-type were derived by taking only peaks which were present in both replicates using BEDtools^107^ intersect (-wa option). For both softwares, consensus peak sets were derived taking all peaks present in one or more cell type using BEDtools intersect (-wa option). Annotation of peaks was performed using ChIPseeker^108^ in R studio. GenomicRanges^109^ was used to measure overlaps and distances between TSSs and peaks.

### ChIP-seq data analysis

Reads were first processed using Trimmomatic to remove adaptors and they were trimmed to 100 bps. They were then processed as above for ATAC-seq except for analysis of transposon coverage (Figure 5B/C) when the reads were not filtered for quality score. Transposons were annotated using RepeatMasker using the default setting and utilizing a previously published manual annotation of *S. rosetta* transposons^70^. Only annotated regions greater than 500 bps were retained. We also only analyzed those families where there were greater than 10 elements present in the genome, thus leaving a group of Mutator-like transposable elements (MULEs) which are DNA transposons and a group of LTR/gypsy-like retrotransposons.

### RNAseq analysis

RNAseq data for the different cell types was published previously^62^. We re-analysed the data by first mapping the reads to the genome using STAR^110^ (--quantMode GeneCounts options). BigWig files were generated using bamCoverage (--binsize 10 –effectiveGenomesize 55000000 – normalizeUsing RPKM) and visualized using IGV. Differential expression analysis was performed using DESeq2^111^. Normalized RPKM values from DESeq2 were used to calculate the log (RPKM+1) values. Visualizations were generated using ggplot2^112^.

### Mass-Spectrometry data analysis

Histone Mass spectrometry data was processed using EpiProfile 2^113^. Briefly, *S. rosetta* histone sequences underwent in silico digestion into peptides, with cleavage occurring after Arginine. Each peptide was then assessed for common potential post-translational modifications (PTMs), such as H3 3-8 unmodified, K4me1, K4me2, K4me3, and K4ac. The area under the curve (AUC) for all peptides is extracted from the raw data. To normalize and facilitate group comparisons, the percentage of each peptide within the same sequence is calculated by dividing its AUC by the summed AUC. The program is available for download at GitHub (https://github.com/zfyuan/EpiProfile2.0_Family/blob/master/EpiProfile2.1_S.rosetta.zip). A summary of the data analysis is available as Supplementary data file 1 which includes AUC, calculated ratios of modified peptides and retention times.

## Data availability

We used the *S. rosetta* genome^58^ (GenBank assembly accession: GCA_000188695.1) available from Ensembl protists (https://protists.ensembl.org/Salpingoeca_rosetta_gca_000188695/Info/Index?db=core) for all analyses. RNA-seq data was published previously^62^ and have been deposited in NCBI’s Gene Expression Omnibus and are accessible through GEO Series accession number GSE267344 (https://www.ncbi.nlm.nih.gov/geo/query/acc.cgi?acc=GSE267344). ATAC-seq and ChIP-seq data generated in this study have been deposited to the NCBI Short Read Archive. ATAC-seq is bioproject PRJNA1107385 and ChIPseq data is bioproject PRJNA1112805,

## Funding

This work was funded by a Wellcome Trust, Sir Henry Wellcome Postdoctoral Fellowship (222767/Z/21/Z) to J.M.G and an NIH award to D.S.B (R35GM147404).

## Contributions

Conceptualization, J.M.G. and D.S.B.; methodology, J.M.G., L.H., L.W., N.V.B., D.S.B.; software, Z.Y.; formal analysis, Z.Y., J.M.G.; investigation, J.M.G., L.H., L.W., N.V.B.; writing, J.M.G. and D.S.B.; supervision, R.K., B.A.C., and D.S.B.; funding acquisition, J.M.G., R.K., and D.S.B. All authors edited and approved the final manuscript.

## Acknowledgements

We would like to thank Nicole King for generously supporting ATAC-seq experiments at an early phase of this project. We would also like to thank Neil Blackledge and the rest of the Klose lab for help and advice with ChIP-seq and analysis and Alex de Mendoza for providing the RepeatMasker annotation. We thank Uri Frank for constructive comments on the manuscript.

## Notes

### Competing Interest Statement

The authors have declared no competing interest.

