## Supplementary materials for "Chromatin profiling identifies putative dual roles for H3K27me3 in regulating transposons and cell type-specific genes in choanoflagellates"

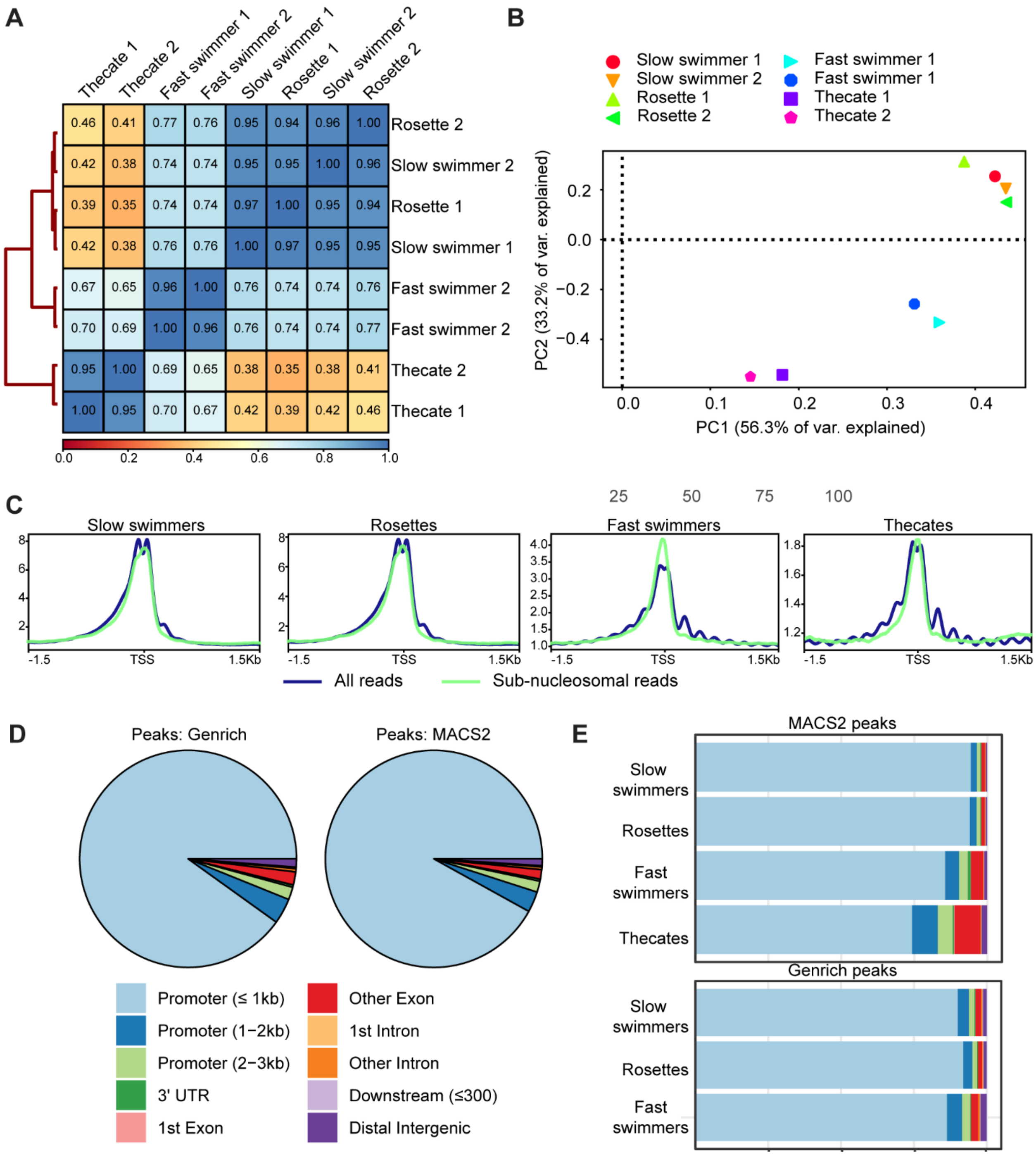


**Figure S1. Additional ATAC-seq analysis**. (**A**) Clustered heatmap showing the correlation coefficients between samples based on the Pearson method. (**B**) PCA plot on all samples showing the two first principal components. (**C**) Metaplot of ATAC-seq read density around the TSS's of all genes comparing all reads to sub-nucleosome sized fragments. **(D)** ChIPseeker annotation of peaks from either Genrich or MACS2. For MACS2 peaks from all cell types are shown while for Genrich all cell types except thecate cells were used (**E**) ChIPseeker annotation of peaks showing each cell type separately.

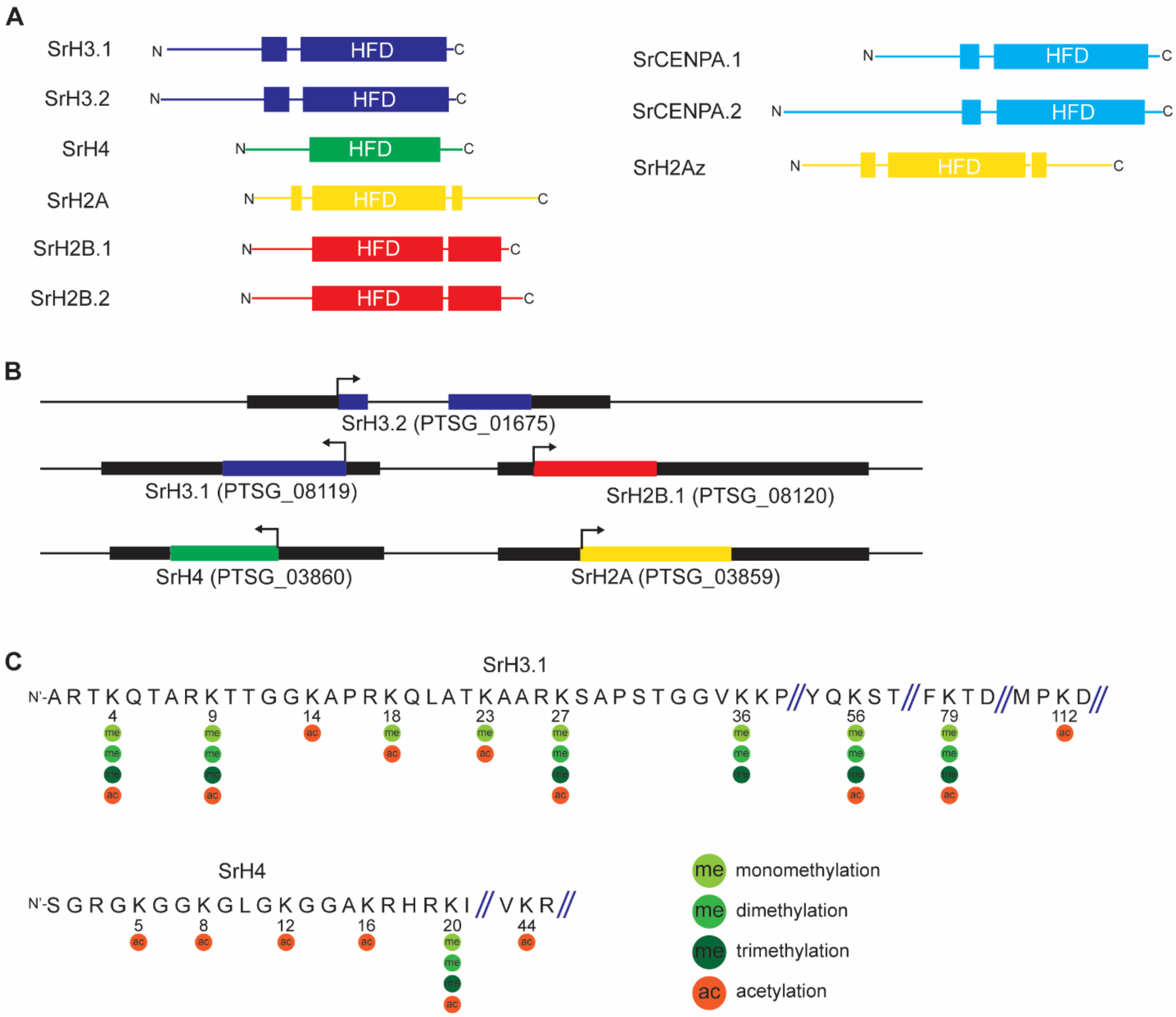


**Figure S2. The histone PTM complement of *S. rosetta*. (A)** Schematic models of all histone proteins annotated in *S. rosetta*. We annotated the histone complement using blast searches as well as previously published phylogenetic analyses (Grau-Bové et al. 2021). S. rosetta has two H3 variants which are putatively annotated as a replication-dependent SrH3.1 and a replication-independent SrH3.2. This is due to SrH3.1 being in a cluster with SrH2A as expected for a canonical replication-dependent histone while SrH3.2 is a single gene which also contains an intron characteristic of a replication-independent variant (see **B**) (Henikoff and Smith 2015). S. rosetta has a single H4 protein, two H2B variants with a variable c-terminal length and a single H2A which is a H2Ax based on the presence of characteristic SQEY amino-acid motif in the C-terminal region. In addition to the canonical histones, *S. rosetta* possesses two centromeric histone H3 variants, SrCENPA.1 and SrCENPA.2 and an H2Az homolog, SrH2Az. (**B**) Genomic organization of the single *SrH3.2* gene and examples of *SrH3.1*, *SrH4*, *SrH2B.1* and *SrH2A*. *Sr.H3.2* is a single copy in the genome and contains an intron. SrH3.1 and SrH2B are in a cluster as are Sr.H4 and SrH2A. (**C**) Schematic of part of the HrH3.1 and SrH4 protein sequence showing positions where lysine methylation and acetylation were identified by mass-spectrometry.

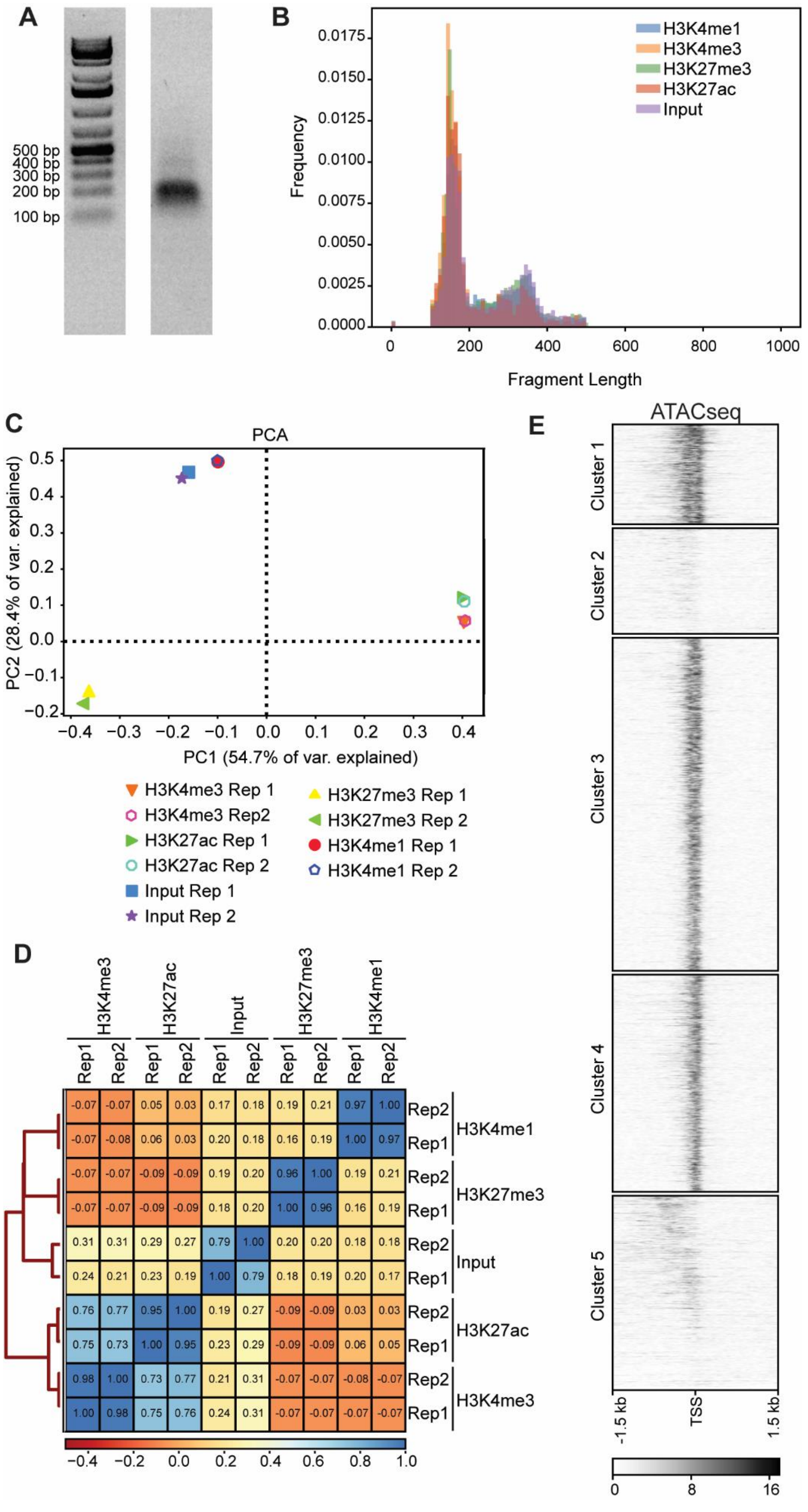
A
500 bp
400 bp
300 bp
200 bp
100 bp
B
H3K4me1
H3K4me3
H3K27me3
H3K27ac
Input
Frequency
0.0000
0.0025
0.0050
0.0075
0.0100
0.0125
0.0150
0.0175
0
200
400
600
800
1000
Fragment Length
C
PCA
PC2 (28.4% of var. explained)
PC1 (54.7% of var. explained)
0.5
0.4
0.3
0.2
0.1
0.0
−0.1
−0.2
−0.4
−0.3
−0.2
−0.1
0.0
0.1
0.2
0.3
0.4
H3K4me3 Rep 1
H3K4me3 Rep2
H3K27ac Rep 1
H3K27ac Rep 2
Input Rep 1
Input Rep 2
H3K27me3 Rep 1
H3K27me3 Rep 2
H3K4me1 Rep 1
H3K4me1 Rep 2
D
H3K4me3
H3K27ac
Input
H3K27me3
H3K4me1
Rep1 Rep2 Rep1 Rep2 Rep1 Rep2 Rep1 Rep2 Rep1 Rep2
-0.07 -0.07 0.05 0.03 0.17 0.18 0.19 0.21 0.97 1.00 Rep2
-0.07 -0.08 0.06 0.03 0.20 0.18 0.16 0.19 1.00 0.97 Rep1
H3K4me1
-0.07 -0.07 -0.09 -0.09 0.19 0.20 0.96 1.00 0.19 0.21 Rep2
-0.07 -0.07 -0.09 -0.09 0.18 0.20 1.00 0.96 0.16 0.19 Rep1
H3K27me3
0.31 0.31 0.29 0.27 0.79 1.00 0.20 0.20 0.18 0.18 Rep2
0.24 0.21 0.23 0.19 1.00 0.79 0.18 0.19 0.20 0.17 Rep1
Input
0.76 0.77 0.95 1.00 0.19 0.27 -0.09 -0.09 0.03 0.03 Rep2
0.75 0.73 1.00 0.95 0.23 0.29 -0.09 -0.09 0.06 0.05 Rep1
H3K27ac
0.98 1.00 0.73 0.77 0.21 0.31 -0.07 -0.07 -0.08 -0.07 Rep2
1.00 0.98 0.75 0.76 0.24 0.31 -0.07 -0.07 -0.07 -0.07 Rep1
H3K4me3
−0.4 −0.2 0.0 0.2 0.4 0.6 0.8 1.0
E
ATACseq
Cluster 1
Cluster 2
Cluster 3
Cluster 4
Cluster 5
-1.5 kb
TSS
1.5 kb
0
8
16

**Figure S3. Additional ChIP-seq data**. (**A**) Gel electrophoresis of DNA from a nucleosome extraction used for ChIP-seq. The majority of fragments are approximately mono-nucleosome size (~140bps). (**B**) Frequency plot showing the size of sequenced fragments from ChIP-seq data. A bin size of 10bp was used. In all 4 antibody ChIPs and input the majority of fragments are below 200bps, as expected for mono-nucleosome fragments. (**C**) PCA plot on all ChIP-seq samples showing the high correlation between replicates. (**D**) Clustered heatmap showing the correlation coefficients between samples based on the Pearson method. (**E**) Heatmap showing ATAC-seq in Slow Swimmers for all genes divided by cluster. Genes are ordered as in Figure 4B. Cluster 1 (n= 1232), Cluster 2 (n= 1305), Cluster 3 (n= 4112), Cluster 4 (n= 2681), Cluster 5 (n= 2401),

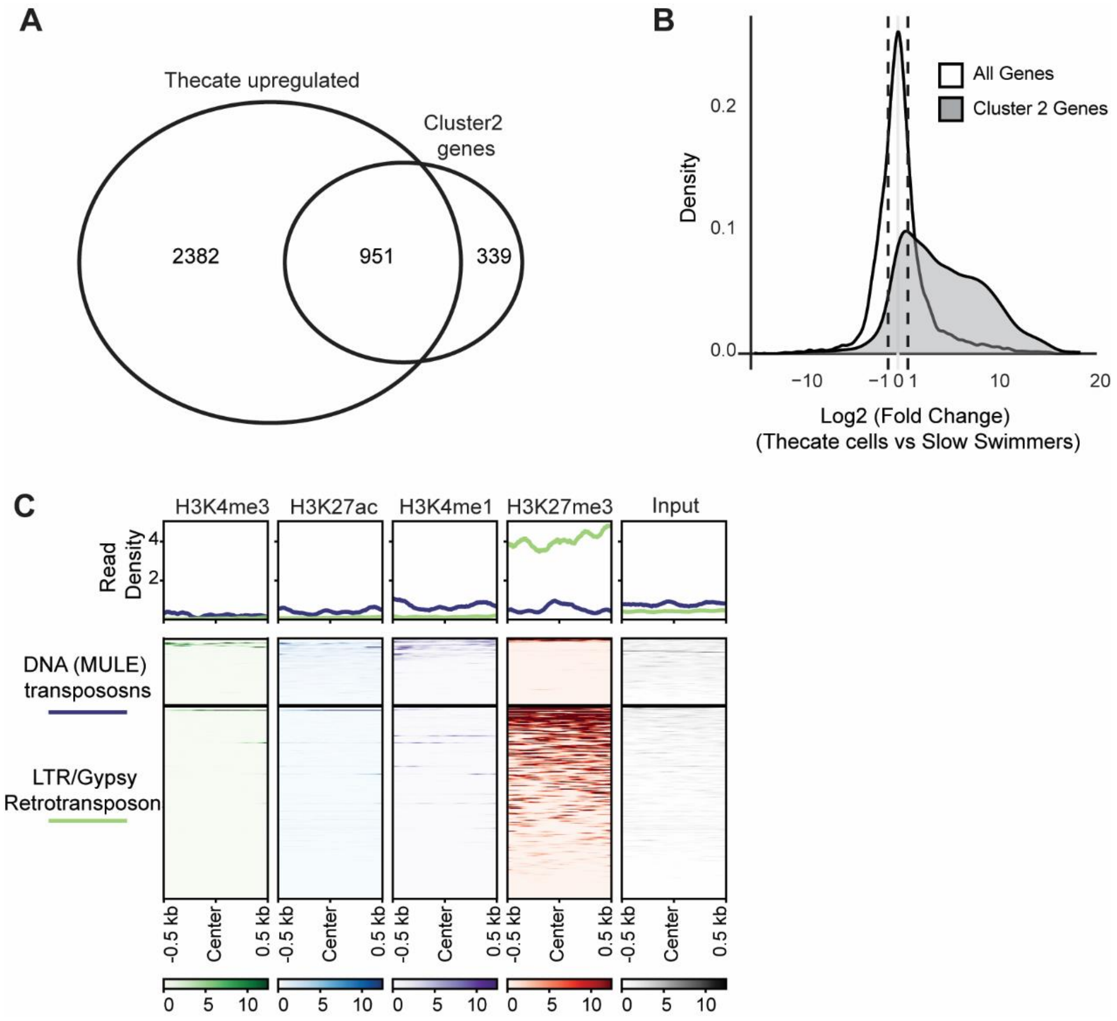


**Figure S4. Additional analysis of H3K27me3 marked regions.** (**A**) Venn diagram showing the overlap between genes from Cluster 2 and genes upregulated in thecate cells vs slow swimmers. Log2 fold change of +/- 1, as determined using DEseq2, was considered as a significant change. (**B**) Density plot showing the log2 fold change between thecate cells and slow swimmers for all genes and for the cluster 2 genes. $Log_2$ fold change was calculated using DESeq2. (**C**) Metaplot and heatmap showing ChIP-seq for the indicated antibodies and input over DNA(MULE) transposons (n=94) and LTR/Gypsy transposons (n=269). Reads here were the same as used in all other analyses (i.e. including base quality cut-off).

**Supplementary Table 1. Histone genes present in *S. rosetta* and their genomic locations.**

| Histone (protein) | Gene names | Genomic location (Supercontig:position) |
|---|---|---|
| SrH3.1 | PTSG_08119 | GL832975:510871-511825 |
| | PTSG_10057 | GL832986:418356-419309 |
| Sr.H3.2 | PTSG_01675 | GL832957:484880-486601 |
| SrH4 | PTSG_03860 | GL832962:1811404-1812225 |
| | PTSG_00674 | GL832956:439532-440249 |
| SrH2A | PTSG_03859 | GL832962:1810216-1811282 |
| | PTSG_00675 | GL832956:440469-441465 |
| SrH2Az | PTSG_11229 | GL832996:49505-51660 |
| SrH2B | PTSG_08120 | GL832975:512187-513213 |
| | PTSG_10056 | GL832986:416738-417364 |
| SrCENPA.1 | PTSG_09298 | GL832982:51979-53872 |
| SrCENPA.2 | PTSG_08448 | GL832977:831645-832995 |

**Supplementary Table 2. List of antibodies used in this study.**

| Target | Source | Concentration used |
|---|---|---|
| H3K27me3 | (Rose et al. 2016) | 5ul per ChIP |
| H3K4me3 | (Farcas et al. 2012) | 5ul per ChIP |
| H3K4me1 | Cell Signalling Technologies D1A9 | 5ul per ChIP |
| H3K27ac | Cell Signalling Technologies D5E4 | 5ul per ChIP |

**Supplementary Table 3. List of software used.**

| Software | Reference | Website |
|---|---|---|
| Illumina BCL Convert v4 | | https://support-docs.illumina.com/SW/BCL_Convert_v4.0/Content/SW/BCLConvert/BCLConvert.htm |
| Bedtools v2.26.0 | (Quinlan and Hall 2010) | https://bedtools.readthedocs.io/en/latest/ |
| Trimmomatic v0.39 | (Bolger et al. 2014) | http://www.usadellab.org/cms/?page=trimmomatic |
| Bowtie2 v2.3.4.1 | (Langmead and Salzberg 2012) | http://bowtie-bio.sourceforge.net/bowtie2/index.shtml |
| Picard v2.18.26 | | https://broadinstitute.github.io/picard/ |
| SAMtools v1.7 | (Li et al. 2009) | https://www.htslib.org/ |
| deepTools v3.4.3 | (Ramírez et al. 2014) | https://deeptools.readthedocs.io/en/develop/ |
| Macs2 v2.1.1 | (Zhang et al. 2008) | https://pypi.org/project/MACS2/ |
| Genrich v0.6.2 | | https://github.com/jsh58/Genrich |

| | | |
|---|---|---|
| STAR v020101 | (Dobin et al. 2013) | https://github.com/alexdobin/STAR |
| ChIPseeker | (Yu et al. 2015) | https://www.bioconductor.org/packages/release/bioc/html/ChIPseeker.html |
| DESeq2 | | https://bioconductor.org/packages/release/bioc/html/DESeq2.html |
| GenomicRanges | (Lawrence et al. 2013) | https://bioconductor.org/packages/release/bioc/html/GenomicRanges.html |
| RepeatMasker | (Smit 2013-2015) | https://www.repeatmasker.org/ |
| IGV | (Robinson et al. 2011) | https://igv.org/ |
| ggplot2 | (Wickham 2016) | https://ggplot2.tidyverse.org/ |
| EpiProfile2.0 | (Yuan et al. 2018) | https://github.com/zfyuan/EpiProfile2.0_Family |
